# Massive detection of cryptic recessive genetic defects in cattle mining millions of life histories

**DOI:** 10.1101/2023.09.22.558782

**Authors:** Florian Besnard, Ana Guintard, Cécile Grohs, Laurence Guzylack-Piriou, Margarita Cano, Clémentine Escouflaire, Chris Hozé, Hélène Leclerc, Thierry Buronfosse, Lucie Dutheil, Jeanlin Jourdain, Anne Barbat, Sébastien Fritz, Marie-Christine Deloche, Aude Remot, Blandine Gaussères, Adèle Clément, Marion Bouchier, Elise Contat, Anne Relun, Vincent Plassard, Julie Rivière, Christine Péchoux, Marthe Vilotte, Camille Eche, Claire Kuchly, Mathieu Charles, Arnaud Boulling, Guillaume Viard, Stéphanie Minéry, Sarah Barbey, Clément Birbes, Coralie Danchin-Burge, Frédéric Launay, Sophie Mattalia, Aurélie Allais-Bonnet, Bérangère Ravary, Yves Millemann, Raphaël Guatteo, Christophe Klopp, Christine Gaspin, Carole Iampietro, Cécile Donnadieu, Denis Milan, Marie-Anne Arcangioli, Mekki Boussaha, Gilles Foucras, Didier Boichard, Aurélien Capitan

## Abstract

We present a data-mining framework designed to detect recessive defects in livestock that have been previously missed due to a lack of specific signs, incomplete penetrance, or incomplete linkage disequilibrium. This approach leverages the massive data generated by genomic selection. Its basic principle is to compare the observed and expected numbers of homozygotes for sliding haplotypes in animals with different life histories. Within three cattle breeds, we report 33 new loci responsible for increased risk of juvenile mortality and present a series of validations based on large-scale genotyping, clinical examination, and functional studies for candidate variants affecting the *NOA1*, *RFC5,* and *ITGB7* genes. In particular, we describe disorders associated with *NOA1* and *RFC5* mutations for the first time in vertebrates. The discovery of these many new defects will help to characterize the genetic basis of inbreeding depression, while their management will improve animal welfare and reduce losses to the industry.

---

Cattle breeds are populations of limited effective size, subject to recurrent outbreaks of recessive defects. Historically, surveillance networks have been established to allow early detection of animals with distinctive clinical features and positional cloning of the locus. With the tremendous development of high-throughput genotyping and sequencing technologies, alternative strategies have been devised to detect genetic conditions that may easily go unnoticed due to a lack of specific signs. These include the search for homozygous haplotype deficiency (VanRaden et al. 2011, Fritz et al. 2013), reverse genetics (e.g. Michot et al. 2016; Charlier et al. 2016) and non-additive association analysis using proxy phenotypes (Reynolds et al. 2021), which have been particularly effective in mapping loci responsible for embryonic or perinatal death and growth retardation. However, these methods are not suitable in cases of incomplete penetrance or incomplete linkage between the causative variant and surrounding markers, which is typically the case for emerging defects due to recent mutations.

To address this issue, we have developed an integrative approach, which is summarized in Fig. 1. The first steps are to list animals genotyped with SNP arrays for routine genomic evaluation, and to mine databases for information on events that shaped their lives to define groups of individuals with distinct life histories (Fig. 1a,b). As a proof of concept, we considered 16,655 heifers that died of natural causes during the rearing period and 488,967 adult cows from the Holstein, Montbeliarde, and Normande breeds (Supplementary Table 1). We analyzed phased Illumina BovineSNP50 genotypes, searching for sliding haplotypes of 20 markers, with at least 25% enrichment in dead heifers and 25% depletion in cows in terms of observed versus expected homozygous animals based on the genotypes of their paternal and maternal ancestors. This approach, which we named “homozygous haplotype enrichment/depletion” (HHED) mapping, yielded numerous candidate regions (Fig. 1c and 2; Supplementary Tables 2,3). For further analysis, we selected the top 20 peak haplotypes per breed and examined the life histories of 8.8 million females born before 2016 whose sires and maternal grandsires (MGS) were genotyped (Fig. 1e,d). After comparing the number of animals in three categories (“dead heifers”, “cows”, and “other”; see Methods) between the offspring of at-risk (carrier sires and carrier MGS) and control mating, we validated 34 of the 60 haplotypes (13, 11, and 10 in the three breeds, respectively) with frequencies ranging from 1.5 to 7.6% (Benjamini-Hochberg adjusted chi-square p-value ≤0.05; Supplementary Tables 4,5). We then analyzed 5 million recent insemination records with sire and dam genotyped. We found 3.4, 3.6 and 8.0 % of at-risk mating in the three breeds and, by extrapolation, estimated that more than 293 thousand calves were born homozygous for at least one locus of increased risk of juvenile mortality in France during the last 10 years (Supplementary Table 6).

**Figure 1.**
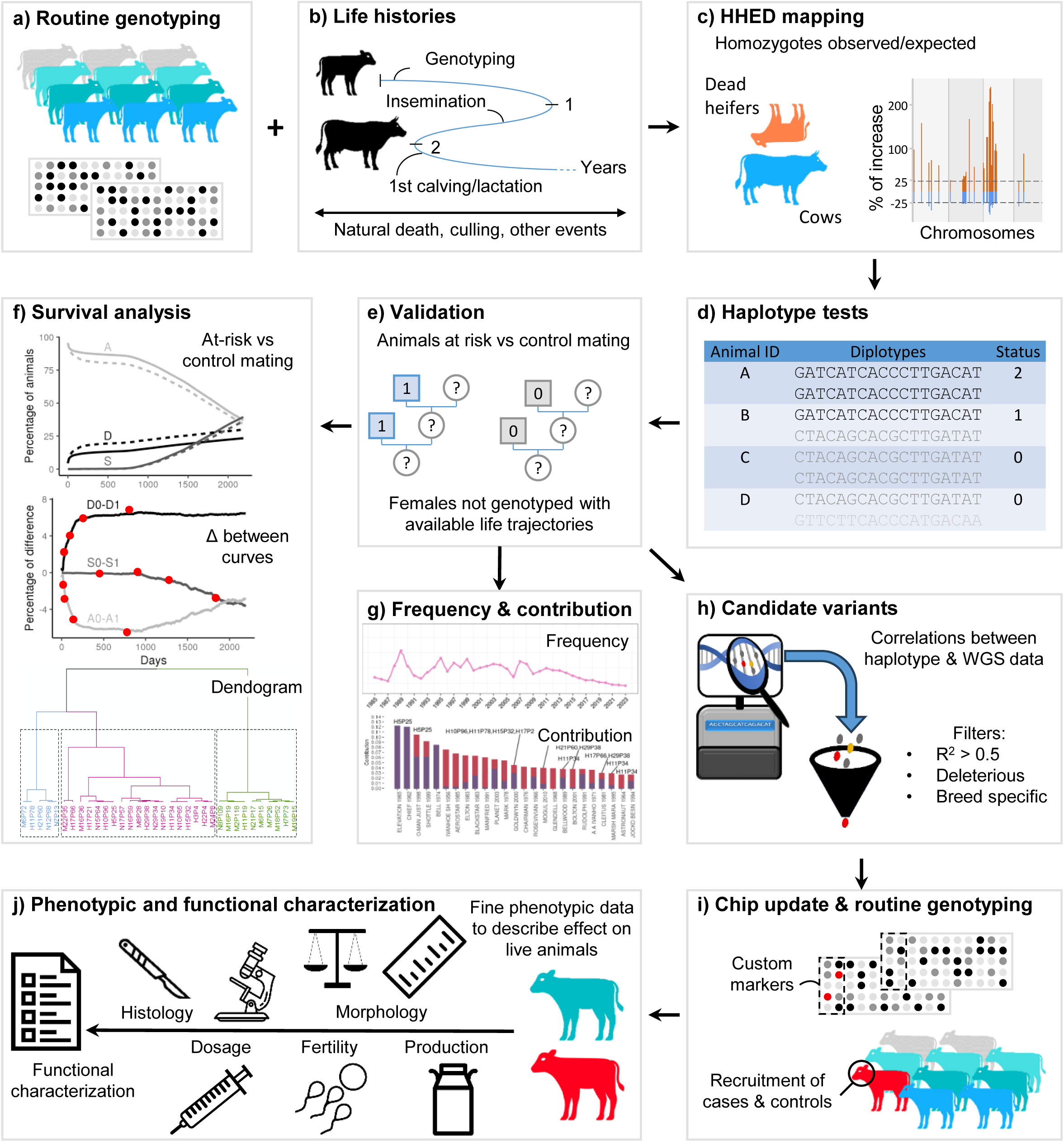
Overview of the analysis plan developed in this study to detect and characterize cryptic recessive genetic defects in cattle. Graphs are shown for illustrative purposes only. Detailed information is provided throughout the manuscript. The first two steps consist of listing animals genotyped for genomic evaluations that have genotypes available for their ancestors (a), and extracting information from national databases to form groups of thousands to tens of thousands of individuals with different life histories (b). Here we focused on juvenile mortality, but the strategy is applicable to other physiological stages and pathologies (premature culling due to infertility or poor production, pre-or post-partum death of dams, etc.). Then phased SNP array genotypes are searched for sliding haplotypes of 20 markers, with at least 25% enrichment in the affected group and 25% depletion in the unaffected group in terms of observed versus expected homozygotes based on the genotypes of their paternal and maternal ancestors (c). Haplotype tests are set up to provide a status (0, 1, or 2 copies) for all genotyped individuals for a selection of the most promising loci (here n=20 per breed; d). The life history of millions of animals born from at-risk or control mating (which are mostly not genotyped) is analyzed and the proportions of animals meeting the inclusion criteria in the affected and unaffected groups are considered for validation (e). These data are also used to characterize survival curves over time using various parameters, principal component analysis, and hierarchical clustering (f). The evolution of haplotype frequencies over time and their causes (e.g. founder effect, heterozygous advantage) are also studied (g). Whole genome sequences from thousands of animals with available haplotype status are then analyzed to identify candidate variants (h). Some of these variants may have been included as markers in the SNP arrays used for genomic evaluation in the framework of reverse genetics studies and have genotypes available for large populations over the last decade (i). These data are used to recruit case and control animals for pathophysiological and functional characterization and to estimate the effects of these variants on survival and a variety of traits at the population level (j). Icons are from www.thenounproject.com.

**Figure 2.**
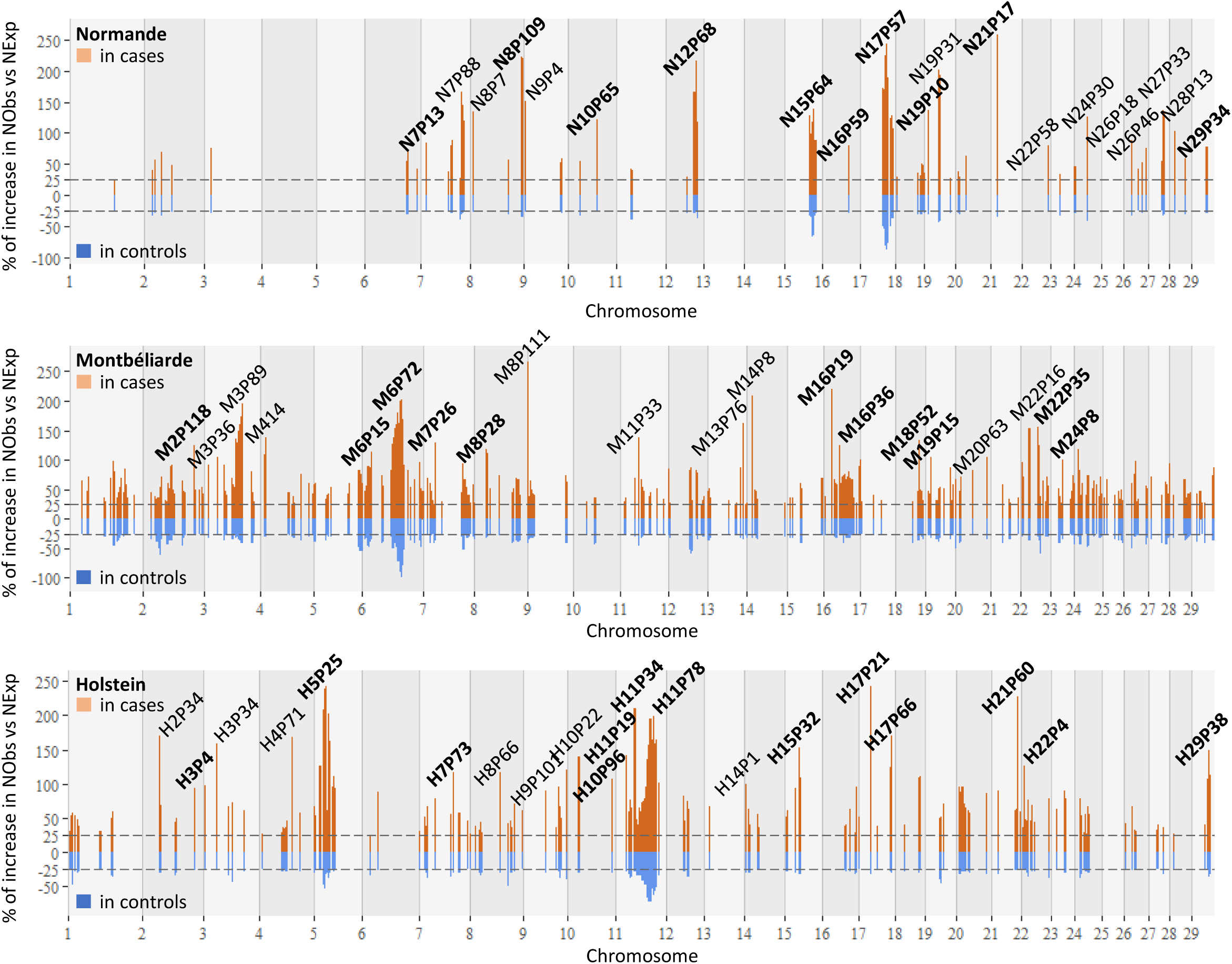
Results of HHED mapping. NObs and NExp: Number of observed and expected homozygous animals, respectively for sliding haplotypes of 20 markers (see Methods). Data supporting the graphs and details of the haplotypes are presented in Supplementary Table 3. The top 20 loci per breed are highlighted (also see Supplementary Table 4), and those validated after analysis of the life histories of 8.8 million females born from at-risk and control mating (see below) are bolded (Benjamini-Hochberg adjusted chi-squared p-value <0.05; Supplementary Table 5). Note that the haplotype names are based on the breed (first character), chromosome (first number), and position (P) in Mb (last number).

To better characterize the effects of the 34 validated haplotypes on female survival, we then calculated the daily difference in proportions between at-risk (1) and control (0) mating for animals that died of natural causes (D), were slaughtered (S), or were still alive (A) over six years (Supplementary Tables 7,8). We then scored the days on which 25, 50, 75, and 100 % of the maximum deviation between the proportion differences were reached (Supplementary Table 9). Using these 12 parameters, we performed a principal component analysis and a hierarchical clustering (Fig. 3a,b), which distinguished three categories of haplotypes according to age and cause of death: i) early juvenile mortality, ii) late juvenile mortality, and iii) increased mortality and premature culling throughout life (Fig. 3c,d, e,f and g,h, respectively). It is worth noting that although the penetrance of juvenile mortality was incomplete for the latter group of loci, we were still able to detect them with our approach. We would also like to highlight that only one haplotype (H11P78) was in linkage disequilibrium with a deleterious variant previously reported in the literature, namely a transposable element insertion in the gene encoding the apolipoprotein B (*APOB*; R^2^=0.66 based on 721,006 genotypes; Supplementary Table 10). APOB-deficient Holstein calves have been reported to suffer from cholesterol deficiency (CD) and die within the first weeks or months of life (Kipp et al. 2016; Menzi et al. 2016; Mock et al. 2016), which is in perfect agreement with our results and provides a validation of our approach (Fig. 3d).

**Figure 3.**
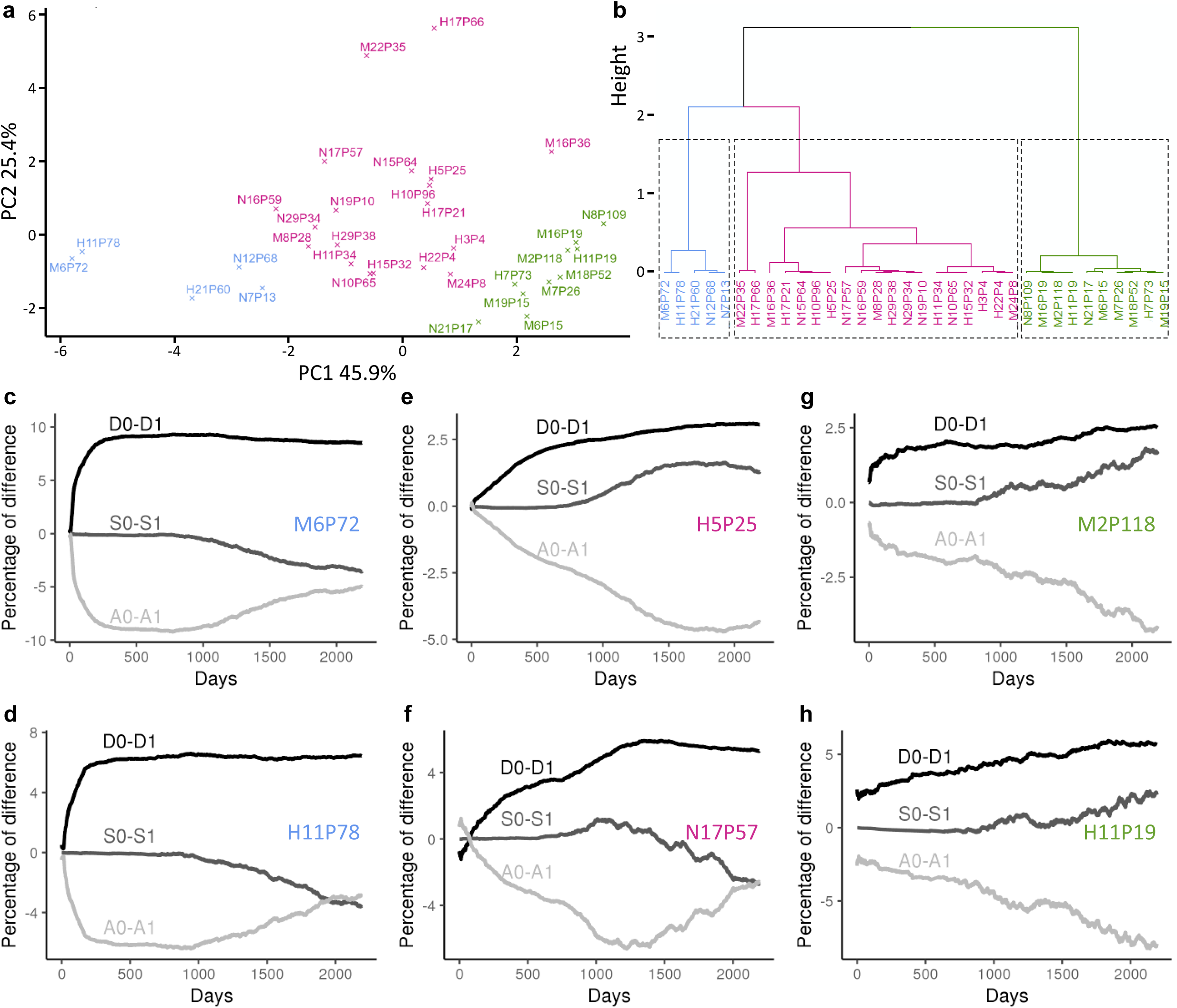
Effects of the 34 validated haplotypes on female survival. a,b) Results of principal component analysis (PCA; a) and hierarchical clustering (HC; b) for 34 haplotypes x 12 parameters (Supplementary Table 9). PC1 and PC2: Proportion of variance explained by the principal components for the first two PCA dimensions. c-h) Daily difference in proportions between at-risk (1) and control (0) mating for animals that died of natural causes (D), were slaughtered (S), or were still alive (A) over six years (Supplementary Table 8). Results are shown for two haplotypes for each of the three PCA-HC clusters (see color code). Note that they include H11P78, which corresponds to the CDH locus (see below), and three other haplotypes that were subjected to clinical and functional investigations later in the article (H5P25, M6P72, and N17P57).

As a next step, we examined the evolution of haplotype frequencies over time (Fig. 4a; Supplementary Tables 11,12). We observed sudden increases in frequency associated with the overuse of influential ancestors, and then slight and regular decreases, most likely due to the massive dissemination of semen from new elite bulls, diluting the genetic contribution of previous ones to the population. The sire GOLDWYN caught our attention, as he is the main source of seven of the 13 haplotypes validated in Holstein cattle, including H11P78/CD (Fig. 4b). This bull was very popular in the late 2000s because of the outstanding functional qualities of his daughters and because he was less related to the population than other champions at that time. This example reminds us of the importance of a more balanced use of breeding stock, as new lines introduce genetic variability, but also potentially new deleterious recessive variants.

**Figure 4.**
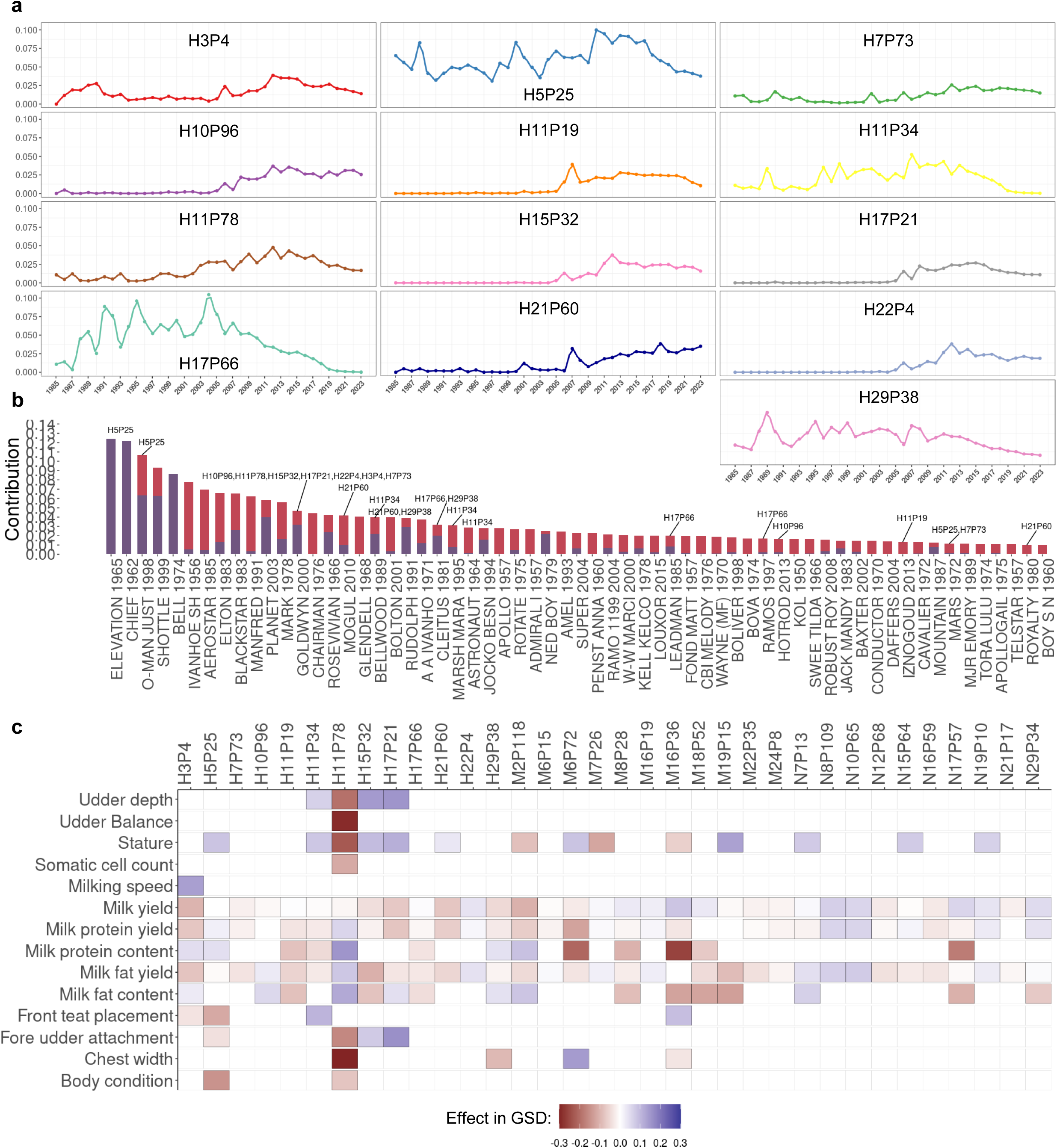
Evolution of haplotype frequencies, pedigree analyses, and estimation of haplotype effects on recorded traits. a) Evolution of frequency for 13 Holstein haplotypes between 1985 and 2023 (Supplementary Table 11). b) Representation of the marginal (purple) and raw genetic contribution of the most used sire (more than 1% of raw contribution) in the French Holstein female population born within the period 2019-2022 (3,058,756 animals) based on pedigree information (see Methods; Supplementary Table 12). c) Effects of the heterozygosity for each Holstein, Montbeliarde, and Normande haplotype on 14 traits expressed in genetic standard deviation (GSD; Supplementary Table 13). Only significant effects are shown (Benjamini-Hochberg adjusted Student’s t-test p-value ≤ 0.01). Note the significant negative effects observed for H11P78/CDH on several traits, including chest width and stature (-0.30 and -0.23 GSD, respectively), which support reduced growth in heterozygous carriers.

In parallel, we estimated the effects of the 34 haplotypes in the heterozygous state on 14 traits routinely collected for selection purposes (see Methods; Supplementary Table 13; Fig. 4c). Due to the high statistical power provided by the large size of our cohorts (1,930≤n≤509,258), we found significant effects in 30% of the analyses (Benjamini-Hochberg adjusted Student’s t-test p-value ≤ 0.01). However, most of these were of small magnitude and we conclude that, overall, the new haplotypes identified in this study do not confer any heterozygous advantage. Interestingly, H11P78 even showed substantial negative effects on several traits (Fig. 4c), echoing previous reports of some CD-clinically affected individuals who were only heterozygous for the *APOB* insertion (Häfliger et al. 2019), and of alterations in lipid metabolism in healthy heterozygotes versus controls (Gross et al. 2016). Therefore, our population-level findings shed new light on CD and support the idea that it is a co-dominantly inherited metabolic disorder.

To gain insight into the molecular mechanisms associated with the 33 novel genetic determinants of increased juvenile mortality, we analyzed the whole genome sequences of 247 Holstein, 160 Montbeliarde, and 118 Normande cattle, as well as 1,344 controls from more than 70 breeds (Supplementary Tables 14,15). By focusing on breed-specific variants that were predicted to be detrimental to protein function and had a genotype-haplotype square correlation (R^2^) of 0.50 or greater, we prioritized 8 candidates (Supplementary Table 16). The identification rate (8/33) may seem somewhat low, and we recognize that the causative variants may affect non-coding regions or segregate in more than one breed, but our goal was to first identify the strongest candidate variants, and then recruit case and control individuals for subsequent phenotypic characterization. As part of reverse genetics studies (e.g., Michot et al., 2016; Bourneuf et al. 2017), six of these candidate variants had been included in the design of the SNP arrays used for genomic evaluation in France, and genotypes were available for 0.4 to 1.8 million animals from 15 breeds.

After verifying that these large-scale genotyping data yielded results consistent with those of whole-genome sequence analysis (Supplementary Table 18), we selected one variant per breed and performed a series of complementary analyses to provide a proof of concept.

The candidate for Holstein haplotype H5P25 is a point mutation of the β7 integrin (ITGB7 p.G375S) affecting a residue conserved among 128 vertebrate orthologs located at the contact interface with the α4 integrin (Fig. 5a; Supplementary Table 19). The α4β7 integrin heterodimer is a major cell adhesion molecule expressed on the surface of CD4 T lymphocytes, that mediates their trafficking from the bloodstream to the digestive tissues (Fig. 5b;Gorfu, Rivera-Nieves, and Ley 2009). This essential process of intestinal immunity could be affected by the ITGB7 substitution, which, based on interaction modeling, is predicted to cause a clash between the two chains and reduce their binding affinity (Fig. 5a). To investigate this hypothesis, we necropsied three homozygous mutant heifers. Although we did not find any specific gross lesions, we observed a complete absence of α4β7^pos^ CD45RO^pos^ memory CD4 T cells in the lamina propria of the jejunum (Fig. 5c, Supplementary Table 20). We also examined 13 case-control pairs of heifers on farms. Homozygous mutants showed stunted growth with a 27% reduction in average daily gain and significant alterations in hematological parameters, such as outstanding WBC and lymphocyte counts (Fig. 5c,d; Supplementary Tables 21,22). Taken together, these results support a deficit in CD4 T cell homing and retention in the gut, as well as other serious immune alterations that require further investigation. For these reasons, we have decided to name this new recessive defect "Bovine Lymphocyte Intestinal Retention Defect" (BLIRD).

**Figure 5.**
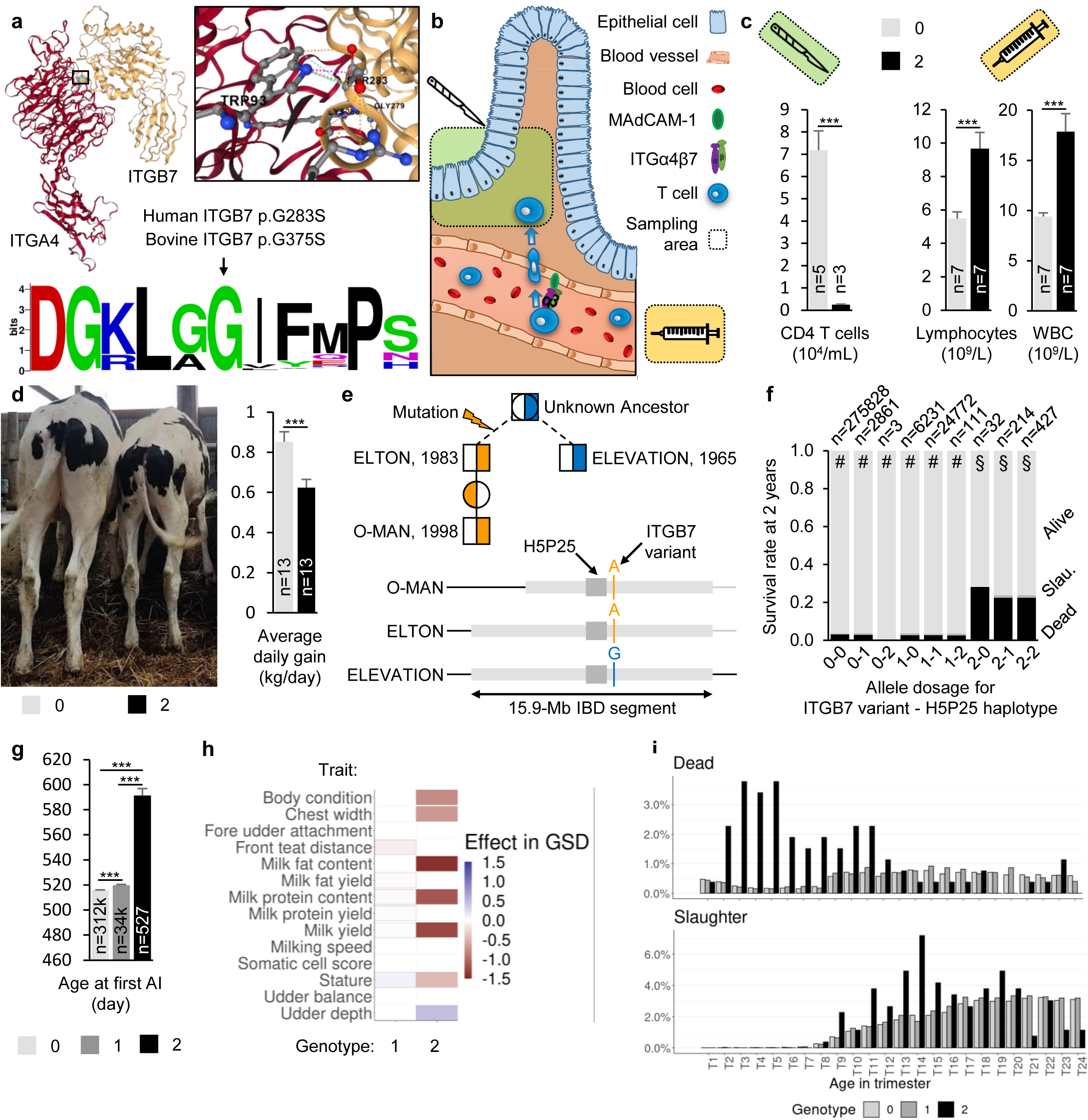
Characterization of the ITGB7 substitution. a top) Modeling of the 3D structure of the ITGα4β7 heterodimer using crystallography in humans, and evaluation of the consequences of glycine to serine substitution at amino acid position 283 in humans (375 in bovine). The purple dashed lines illustrate the clash with a predicted 0.8 kcal/mol reduction in binding affinity (see Methods for further details). a, down) Weblogo representation of multiple protein alignments showing perfect conservation of the mutated glycine residue among 128 vertebrate ITGB7 orthologs (Supplementary Table 19). b) Schematic representation of the ITGα4β7-mediated transmigration of T lymphocytes (T cells) from blood vessels to the lamina propria of the jejunum, and details of the sampling area for subsequent analyses. c) Dosage of α4β7pos CD45ROpos memory CD4 T cells in the lamina propria of the jejunum and of immune cells in the circulating blood of homozygous wild-type (0) and mutant (2) heifers (Supplementary Tables 20,21). ***: Student t test p-value ≤0.001. d) Pictures of 247-day-old wild-type and 245-day-old homozygous mutant heifers, illustrating growth retardation in the latter; and analysis of average daily gain in 13 cases and 13 matched control heifers aged six months to two years (Supplementary Table 22). e) Origin of H5P25 haplotype and *ITGB7* mutation (see Supplementary Table 23 for details). f) Proportion of animals that died, were slaughtered, or are still alive at 2 years of age for each combination of *ITGB7* variant genotype and H5P25 status. Different symbols indicate significant differences (Fisher p-value ≤0.01) between genotype-haplotype combinations in the proportions of animals within each of the three categories (Supplementary Table 24). g) Analysis of age at first AI for the three genotypes at the *ITGB7* variant (Supplementary Table 25). ***: Student t test p-value ≤0.001. Effects of heterozygosity and homozygosity at the *ITGB7* variant on 14 recorded traits (Supplementary Table 26). Only significant effects are shown (Benjamini-Hochberg adjusted Student’s t-test p-value ≤ 0.01). i) Trimester repartition of the proportion of females that died or were slaughtered before reaching six years of age by genotype group at the *ITGB7* substitution (Supplementary Table 27).

Based on pedigree and genotype information, we determined that ELEVATION, the most influential ancestor of the Holstein breed, and ELTON, the grandsire of the third most influential ancestor, O-MAN, shared an identical-by-descent segment of 15.9 Mb encompassing the H5P25 haplotype and that the *ITGB7* mutation occurred in an ancestor of ELTON (Fig. 5e; Supplementary Table 23). Taking advantage of this incomplete LD, we examined the two-year survival of the different haplotype x genotype combinations (see Methods) and observed an excess of mortality only in homozygous mutants, regardless of their haplotype status, further confirming the causality of the ITGB7 substitution (Fig. 5f; Supplementary Table 24). Finally, population-level analyses revealed delayed age at first insemination (Fig. 5g; Supplementary Table 25), consistent with reduced growth (e.g., Jourdain et al. 2023), adverse effects on production traits (Fig. 5h; Supplementary Table 26), and premature culling in homozygous mutants compared to other genotypes (Fig. 5i; Supplementary Table 27). As a recessive defect that (i) causes juvenile mortality with incomplete penetrance, (ii) is due to a mutation in incomplete LD with surrounding markers, and (iii) has not been previously detected despite segregating for more than 40 years in Holstein cattle, BLIRD provides a textbook example to validate our approach.

We then studied two variants affecting proteins for which no living homozygous mutants have been described so far in animals: an inframe deletion of the Replication Factor C Subunit 5 affecting a residue conserved in 633 eukaryotic orthologs (RFC5 p.E369del; R^2^=1 with haplotype N17P57; Fig. 6a; Supplementary Tables 16;28), and a frameshift insertion in the Nitric Oxide-Associated Protein 1 (NOA1 p.D400RfsX9; R^2^=0.55 with M6P72; Fig. 6i; Supplementary Table 16).

**Figure 6.**
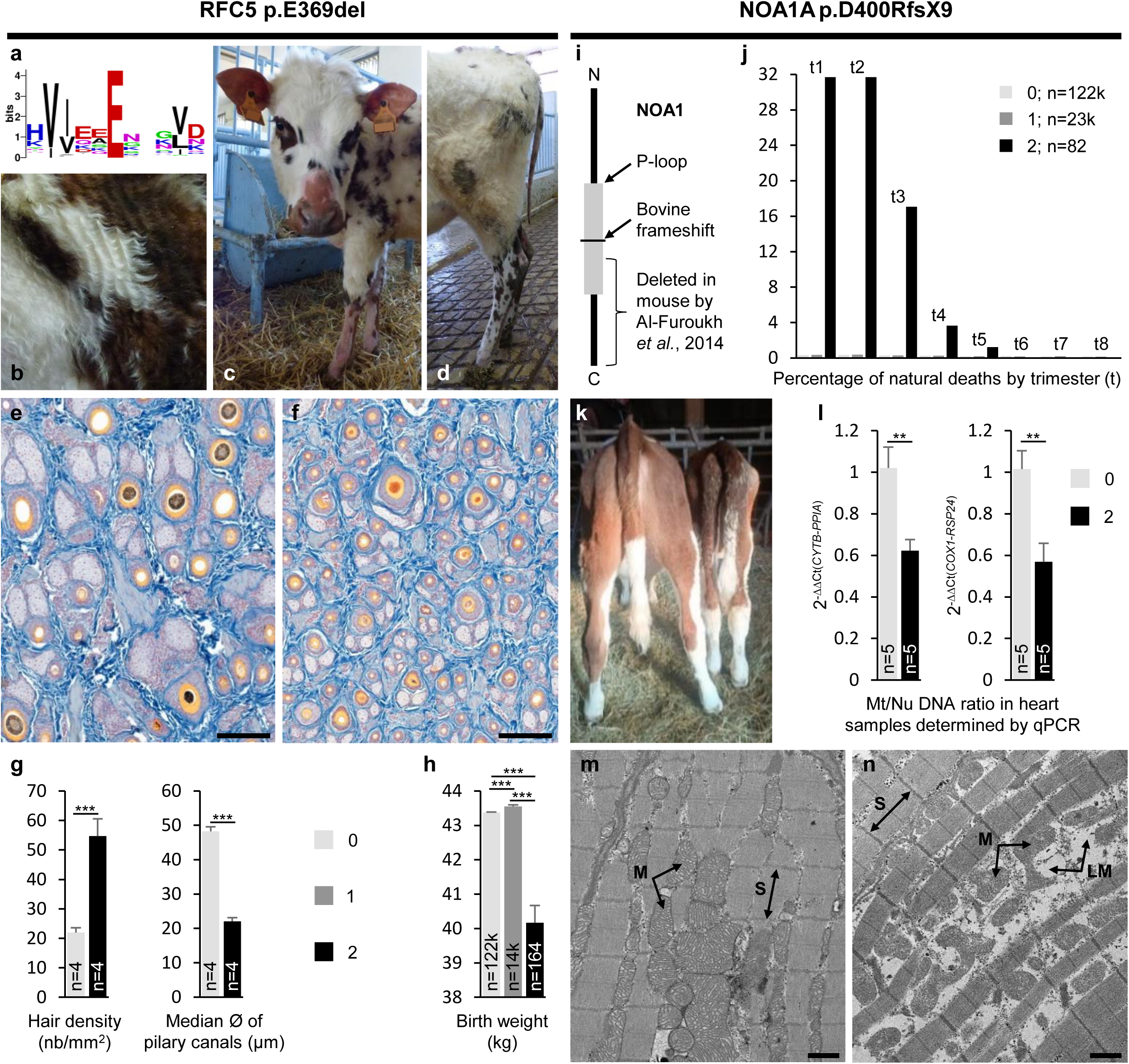
Pathophysiological characterization of RFC5 and NOA1 mutants in Normande and Montbeliarde cattle, respectively. a) Weblogo representation of multiple protein alignments showing perfect conservation of the glutamic acid residue at position 369 amino acids in bovine among 633 eukaryotic RFC5 orthologs (Supplementary Table 28). b) Detail of the shoulder and base of the neck of a homozygous mutant heifer showing a thin and wavy hair coat. c,d) Images of the same animal showing alopecia of the ears, the top of the snout, extremities of the limbs, and tail. e,f) Skin sections of shoulder skin samples from wild-type (e) and *RFC5* homozygous mutant (f) 1.5-year-old heifers stained with a Roan solution. Scale bars = 200 µM. g) Analysis of shoulder skin sections from 4 cases and 4 control heifers for hair density in a randomly selected square of 1 mm^2^, and median size of 50 adjacent pilary canals (Supplementary Table 29). h) Analysis of birth weight for the three genotypes at the *RFC5* deletion (Supplementary Table 30). i) Schematic representation of the bovine NOA1 protein (Ensembl ID: ENSBTAP00000025792, 731 aa). If expressed, the p.D400RfsX9 frameshifted bovine protein would lack half of the P-loop containing nucleoside triphosphate hydrolase domain (P-loop) and a conserved C-terminal region, the deletion of which causes impaired mitochondrial import of NOA1 in cultured mouse myoblasts (Al-Furoukh et al. 2014). j) Percentage of natural deaths by trimester for 145 thousand females genotyped for the *NOA1* frameshift variant. Note that only the first two years of a six-year study are shown (Supplementary Table 35). k) Wild-type (left) and *NOA1* homozygous mutant (right) 4.5-month-old heifers. l) Relative changes in mitochondrial-to-nuclear (Mt/Nu) DNA ratio in heart samples determined by qPCR (Supplementary Tables 37,38). m,n) Transmission electron microscopy images of myocardial samples from wild-type (m) and *NOA1* homozygous mutant (n) four-month-old heifers. Scale bars = 1 µM. S: Sarcomere, the basic unit of myofibrils. M: Mitochondria. LM: Lysed mitochondria. Note the presence of empty spaces between myofibrils due to increased mitochondrial cell death in (n). ** and ***: Student t test p-value ≤0.01 and ≤0.001, respectively. 0, 1 and 2: wild-type, heterozygous and homozygous genotypes for the *RFC5* inframe deletion or *NOA1* frameshift insertion, respectively.

RFC5 is one of the five subunits of the replication factor C complex, which is essential for DNA replication and cell proliferation in eukaryotes (Cullmann et al. 1995; Furukawa et al. 2003; Li et al. 2018). Clinical examination of six homozygous mutant heifers revealed stunted growth, chronic diarrhea with no gross lesions at necropsy, abnormally thin and wavy hair coat, and alopecia of the body extremities (Fig. 6b-d). Breeders reported that hairless patches appeared after the juvenile molt and peaked in size each winter, suggesting that low temperatures impair hair growth in homozygotes. This observation is consistent with the temperature sensitivity reported for two RFC subunit mutants in yeast, which is associated with blockage of DNA replication (N. Reynolds, Fantes, and MacNeill 1999). Histological analysis of shoulder skin samples showed increased hair density and decreased hair diameter in case versus control heifers, demonstrating that the homozygosity at the *RFC5* inframe deletion also affects hair follicle differentiation during embryogenesis (Fig. 6e-g, Supplementary Table 29). Finally, phenotypic characterization at the population level revealed reduced birth weight in homozygous mutants supporting *in-utero* growth retardation, and elevated rates of late juvenile mortality and premature culling (Fig. 6h; Supplementary Tables 30,31; Supplementary Fig. 1).

NOA1 is a nuclear-encoded protein required for mitochondrial protein translation and respiration, and the p.D400RfsX9 mutation is predicted to alter its function and mitochondrial import (Kolanczyk et al. 2011; Al-Furoukh et al. 2014; Fig. 6i). While *NOA1* knockout in mice causes midgestation lethality (Kolanczyk et al. 2011), we estimated that only 24.4% of the homozygous mutants died during pregnancy (Supplementary Table 32). An additional 50.7% died before reaching the age of genotyping (Supplementary Table 33), and the 24.9% that were genotyped died in most cases before one year of age (Supplementary Tables 34,35; Fig. 6j). Thus, the eight two-to four-month-old genotyped homozygotes (one male, seven females) that we followed clinically were among the longest-lived. The females developed ill-thrift between three and 12 months of age and were euthanatized for ethical reasons. Hematological and immune analyses revealed neutrophilia, indicating the presence of inflammation but no anomaly of reactive oxygen species (ROS) production by neutrophils (Supplementary Table 36, Supplementary Note). In addition, *NOA1* mutants showed abnormal blood biochemical parameters suggesting a metabolic disorder and extensive mitochondrial apoptosis, as revealed by electron microscopy and relative quantification of mitochondrial and nuclear DNA in myocardial samples (Supplementary Tables 36-38; Fig. 6l-n). Finally, the only homozygous male never showed clinical signs until the end of its follow-up at one year. In contrast to mice, the distribution of deaths over a long period and access to large populations whose genetic background and rearing conditions are not standardized offer interesting prospects for the future identification of genetic or environmental factors that might compensate for NOA1 loss-of-function in cattle.

In conclusion, using a data science-based approach, we have identified numerous recessive loci responsible for increased risk of juvenile mortality in cattle that had previously been overlooked. The data-mining framework described in this paper is readily applicable to other physiological stages and any population that benefits from large datasets generated by genomic evaluation. The management of these new genetic defects will have a direct impact on animal breeding, helping to reduce animal suffering and economic loss to the industry. Finally, our approach also offers exciting prospects for basic research, by identifying large animal models for immune or metabolic disorders, some of which involve understudied genes, that can be characterized at a population level.

## Methods

### Animals and data sets

A large number of animal populations and data sets were considered in this study. These are detailed below for each analysis. Briefly, the most important of these are: (i) pedigree and life history information on millions of female cattle (date of birth; lifespan; cause of death; dates of insemination and bull IDs; dates of calving and onset of lactation) extracted from the French national bovine database; (ii) phased and imputed Illumina BovineSNP50 array genotypes generated as part of the routine bovine genomic evaluation; (iii) performance for various traits corrected for non-genetic factors as estimated in the national genomic evaluation; (iv) whole genome sequences of 1869 cattle; (v) SNP array genotypes for candidate variants in large populations; and (vi) animals recruited from commercial farms for physiopathological characterization and functional analyses.

### Mapping of recessive loci, validation, and analysis of survival curves

Pedigree and life history information was extracted from the French national bovine database for 5.96 million Holstein, 1.63 million Montbéliarde, and 1.24 million Normande females whose sires and maternal grandsires (MGS) were genotyped. The discovery population was restricted to genotyped females and included 8,203, 6,198 and 2,254 “dead heifers” (females that died of natural causes before 3 years of age and were never inseminated) and 291,529, 141,343 and 56,095 cows (females that calved and started a first lactation) from the three breeds, respectively (Supplementary Table 1). These animals and their ancestors were genotyped using various Illumina SNP arrays over time (LD,∼7 K SNPs; custom LD,∼10 K to 20 K; BovineSNP50,∼50 K; EuroGMD,∼63 K; and HD,∼777K). Raw genotypes were imputed and phased for 44,596 autosomal SNPs by FImpute (Sargolzaei et al. 2014) as part of the French routine genomic evaluation of cattle, as described by Mesbah-Uddin et al. 2019; Supplementary Table 2). We considered sliding haplotypes of 20 markers, and counted the number of homozygotes observed (NObs) within each group of genotyped individuals. In parallel, we estimated the expected number of homozygotes (NExp) using within-family transmission probability. We filtered haplotypes satisfying the following criteria: Nobs ≥10 in cases, increase in homozygotes (i.e. (Nobs-Nexp)/Nexp)) ≥ 25% in cases and ≤ -25% in controls (Supplementary Table 3). Among stretches of consecutive haplotypes, we selected the one showing the highest increase in homozygotes in cases as the “peak haplotype” (Supplementary Table 4). For validation, we compared the proportions of animals belonging to three categories (“dead heifers”, “cows”, and “others”) using a chi-squared test with Benjamini-Hochberg correction (p-value ≤ 0.05) among the descendants of at-risk mating (“1”; carrier sire and carrier MGS) or control mating (“0”; noncarrier sire and MGS). Note that the “other” category includes females that do not meet the criteria retained for the “dead heifers” and “cows” groups (e.g. heifers that were slaughtered, heifers that died after 3 years of age, etc.). This analysis was performed on a validation population of 5.96 million Holstein, 1.63 million Montbeliarde and 1.24 million Normande individuals born between 2000 and 2015, most of whom were not genotyped themselves (Supplementary Table 5). Then, for 34 validated haplotypes, we calculated the daily proportion of animals that died of a natural causes (D), were slaughtered (S) or were still alive (A) over a period of 6 years for mating types 0 and 1 (Supplementary Table 7). We also calculated the D0-D1, S0-S1 and A0-A1 differences in proportion on a daily basis and scored the days on which 25, 50, 75 and 100 % of the maximum deviation between each proportion difference was reached (Supplementary Table 8). We then used these 12 parameters to perform a principal component analysis and a hierarchical clustering using the RStudio package Factoshiny v.1.2.5033 (Supplementary Table 9).

### Estimation of the number of calves born homozygous for one or more validated haplotypes in the last ten years

We considered 55.5 million inseminations performed during the period 2013-2022, of which 2.0 million involved females and males that were both genotyped. To estimate the number of homozygous calves (NH), we took into account the total number of inseminations (NAI), the proportion of at-risk mating observed within genotyped couples (PR), the Mendelian probability (0.25), the average conception rate in each breed (CR), and finally the proportion of females bred by AI (%AI): NH = NAI*PR *CR*0.25/ %AI (Supplementary Table 6; Escouflaire and Capitan 2021.

### Evolution of haplotype frequencies over time

The frequency of the 34 validated haplotypes was calculated on an annual basis considering 1,185,446 Holstein, 591,294 Montbeliarde and 180,722 Normande individuals of any sex available in the French bovine national genomic evaluation database born between 1985 and 2023 (Supplementary Table 11).

### Genetic contribution of the ancestors of the actual female populations

To identify the main ancestors of the current female populations and calculate their raw and marginal genetic contributions, we analyzed the pedigrees of 3,058,756 Holstein, 738,333 Montbeliarde and 333,793 Normande females born within the period 2019-2022 with at least sire and dam information available (Supplementary Table 12) using the PEDIG software (Boichard 2002).

### Effects of haplotypes on recorded traits

The effects of the 34 haplotypes in the heterozygous state were estimated on 14 morphological and production traits routinely recorded for selection purposes. To remove the various environmental factors affecting these phenotypes, we used yield deviation data, i.e. records adjusted for the non-genetic effects included in the genomic evaluation models. Yield deviations are a by-product of the official genomic evaluations carried out by GenEval on behalf of the French breeding organizations (for details on the models used, see https://www.geneval.fr/english). Yield deviations were analyzed with a mixed model including the fixed effect of the haplotype studied (0 versus 1 copy), the fixed effect of the year of recording, and the individual random polygenic effect. Calculations were performed with BLUPF90 software (Misztal et al. 2014). The sample size ranged from 1930 to 509,258 cows per group (Supplementary Table 13). Student’s t-test was used to compare the means between groups for each trait and adjusted using the Benjamini-Hochberg method. To allow comparisons between traits, effects were converted to genetic standard deviations (GSD) based on the genetic parameters estimated for the national genomic evaluations.

Estimating effects in homozygotes did not seem relevant to us because at this stage we cannot know whether the causal variants are in complete linkage disequilibrium with the causative variant. These effects may be strongly biased if, among animals homozygous for the haplotype with performance records, there is a large increase in the proportion of animals that are not homozygous for the causative variant as a result of natural counterselection of double homozygotes (for both the haplotype and the causative variant).

### DNA extraction

Genomic DNA was extracted from semen straws, blood or myocardium samples using the DNeasy Blood and Tissue Kit (Qiagen). DNA quality was controlled by electrophoresis and quantified using a Nanodrop spectrophotometer.

### Whole genome sequencing and variant calling

We analyzed the whole genome sequences of 1,869 cattle from more than 70 different breeds generated by Illumina technology (Supplementary Table 14). Of these, 1,137 were obtained from public databases and 732, consisting mainly of influential AI bulls of French breeds, were sequenced at the GeT-PlaGe facility during the last decade (http://genomique.genotoul.fr/). Paired-end libraries with insert sizes ranging from 300 to 500 bp were generated using the NEXTflex PCR-Free DNA Sequencing Kit (Biooscientific) or the TruSeq Nano DNA Library Prep Kit (Illumina). Libraries were quantified using the KAPA Library Quantification Kit (Cliniscience), checked using a High Sensitivity DNA Chip (Agilent) or the Fragment Analyzer (Agilent), and sequenced on Illumina machines (HiSeq2500, Hiseq3000, Hiseq10X, or NovaSeq 6000) with 2 × 100-bp or 2 × 150-bp read lengths, according to the manufacturer’s protocol. Reads were aligned to the bovine ARS-UCD1.2 reference genome sequence using the Burrows–Wheeler aligner (BWA-v0.6.1-r104; H. Li and Durbin 2009). SNPs and small InDels were identified using the GATK-HaplotypeCaller software (McKenna et al. 2010) as previously described (Daetwyler et al. 2014; Boussaha et al. 2016), while structural variations (SVs) were detected using the Pindel (Ye et al. 2009), Delly (Rausch et al. 2012), and Lumpy (Layer et al. 2014) software.

### Variant filtering and annotation

R^2^-correlations between haplotype status and genotypes for variants located in a 20 Mb region centered on each of the 33 haplotypes were calculated for 247 Holstein, 160 Montbeliarde, and 118 Normande WGS. The number of haplotype carrier alleles ranged from one to 22 in this data set (Supplementary Table 15). Variants with an R^2^ score≥0.5 were selected, and those segregating in more than one breed represented in the full set of WGS were filtered out, assuming that the causative mutations occurred after the creation of the breeds. The remaining variants (consisting only of SNPs and small InDels) were annotated using Variant Effect Predictor (Ensembl release 110; https://www.ensembl.org/info/docs/tools/vep/index.html), and only those predicted to be deleterious were considered (i.e., missense variants with a SIFT score ≤0.05, stop gain or loss, premature start, start loss, frameshift, inframe insertion or deletion, and variants affecting splice donor or acceptor sites). Finally, the consistency of these annotations was verified using independent resources: some variants predicted by VEP to be deleterious and located in non-coding regions (i.e., with no evidence of bovine expressed sequence tag or orthologous transcripts in mouse or human) according to the USC genome browser (http://genome.ucsc.edu/) were further removed. Information on the pathophysiological features associated with mutations affecting orthologous genes in humans and mice was extracted from the Online Mendelian Inheritance in Man (OMIM; https://www.omim.org) and Mouse Genome Informatics (MGI; https://www.informatics.jax.org) databases (Supplementary Table 16). The putative amino acid sequence of the mutant NOA1 protein was obtained using the Expasy translate tool (https://web.expasy.org/translate/) after insertion of the frameshift allele into the cDNA of the Ensembl transcript ENSBTAT00000025792. Information on "domains & features" was obtained from the eponymous Ensembl “transcript-based display”. To analyze amino acid conservation at the ITGB7 and RFC5 mutation sites, “1-to-1” orthologous proteins from numerous animal species were extracted from the “orthologues” “gene-based display” available for Ensembl entries ENSBTAG00000018993 and ENSBTAG00000007137 and aligned using ClustalW software version 2.1 (Thompson, Higgins, and Gibson 1994; https://www.genome.jp/tools-bin/clustalw). Finally, a sequence logo was generated using WebLogo (Crooks et al. 2004; http://weblogo.berkeley.edu). Note that RFC5 in animals is orthologous to RFC3 in fungi and plants, as supported by orthology comparisons with the yeast (S. cerevisiae) protein entry YNL290W that are available in Ensembl release 110 (www.ensembl.org/), Ensembl Fungi release 57 (http://fungi.ensembl.org/), and Ensembl Plants release 57 (http://plants.ensembl.org/index.html). “1-to-1” orthologs of yeast RFC3 were extracted from the latter two databases and added to the animal orthologs of the bovine RFC5 protein for analysis. Information on protein IDs, species, and amino acid sequence around the mutation sites is provided in Supplementary Tables 19 and 28.

### Analysis of the 2/3D structure of the mutant ITGB7 protein

The effect of the bovine p.G375S point mutation on the ITGB7 protein structure was evaluated using the mCSM-PPI2 server (https://biosig.lab.uq.edu.au/mcsm_ppi2/; Rodrigues et al. 2019). For this purpose, the crystal structure of the human ITGA4/ITGB7 dimer interacting with the Act-1 mAb was used as a reference structure model (accession number 3V4P in the Worldwide Protein Data Bank; www.wwpdb.org). The orthologous glycine residue is located at position 283 in the human ITGB7 protein used for modeling.

### Large-scale genotyping of candidate variants and analysis of LD with haplotypes

The Illumina SNP arrays used for genomic evaluation in France have a custom section to which probes for genotyping thousands of deleterious variants have been added over time through various forward and reverse genetic projects. Genotypes were available for the APOB insertion responsible for recessive CD in Holstein cattle and for six candidate variants for new loci associated with increased juvenile mortality discovered in this study. Information on the probes and the number of genotypes available in 15 breeds can be found in Supplementary Tables 10, 16, 17 and 18. These data were used to calculate allele frequencies, to verify the breed specificity of the markers, to compute contingency tables between haplotype status and genotypes, and to calculate R^2^ square correlations. Finally, they were also used to investigate the effect of the ITGB7, RFC5, and NOA1 candidate variants on various traits (see related sections).

### Cause of incomplete LD between some haplotypes and their candidate variants

To determine whether the incomplete LD was due to a relatively recent de novo mutation or to an ancient founder effect followed by recombination events between the haplotype and the variant, we sorted carriers of (i) both the haplotype and the variant, (ii) the haplotype but not the variant, and (iii) the variant but not the haplotype. We searched for IBD segments between these animals and their ancestors using both genotype and pedigree information. Illumina BovineSNP50 array genotypes were available for most of the AI bulls used in France since 1985. This analysis was performed for H5P25/*ITGB7* (Supplementary Table 23) and M6P72/*NOA1* (Supplementary Fig. 2)

### Imputation of *ITGB7* and *RFC5* variants using long size haplotypes

As the *ITGB7* and *RFC5* candidate variants were included in the arrays used for genomic evaluations in France in early 2019, we have developed an approach to impute them in animals genotyped before that date, in order to allow survival studies over 6 years and to increase cohort sizes for studying different traits. For the *ITGB7* substitution, we considered a 34 Mb segment (476 markers from markers ARS-BFGL-NGS-69702 to ARS-BFGL-NGS-7850) centered on the 15.9 IBD segment shared by the bulls ELEVATION and ELTON, which were (i) heterozygous carrier of the H5P25 haplotype but wild type for the variant for the former and (ii) double heterozygous for the latter (Figure 5; Supplementary Table 23). For the *RFC5* inframe deletion, which was in complete LD with N17P57, we arbitrarily considered a 5 Mb segment centered on the inframe deletion (105 markers from Hapmap34428- BES2_Contig387_701 to BTB-00682446). See Supplementary Table 2 for information on the marker map. Using 272,326 Holstein cattle genotyped for the *ITGB7* variant and 53,263 Normande cattle genotyped for the *RFC5* variant, we created a bank of long-size haplotypes associated with either the ancestral or the mutant allele. If a long-size haplotype was not associated with only one allele, it was considered dubious and eliminated. We then proceeded to genotype imputation for 789,594 and 127,783 animals, respectively. Animals with one or two long-size haplotypes unknown in the haplotype bank were not considered. This procedure has been designed to reduce imputation errors to a level close to zero. Finally, we generated a database of 585,671 *ITGB7* genotypes in Holstein and 164,291 *RFC5* genotypes in Normande cattle with 1.15 and 2.08 ratios of imputed/real genotypes, respectively. Imputation was not required for the NOA1 frameshift insertion as it has been genotyped since 2013.

### Analysis of survival based on genotype for candidate

To verify the causality of the *ITGB7* and *NOA1* variants, a first analysis of the proportions of animals still alive at 2 years of age or that died or were slaughtered before was performed for each combination of variant genotype x haplotype status (Supplementary Tables 24,34). This analysis was not performed for the *RFC5* inframe deletion because of its complete LD with the N17P57 haplotype. In addition, for the three genotypes of each variant, we calculated the proportions of animals still alive at 6 years of age or that died or were slaughtered before, and their repartition per trimester (Supplementary Tables 27,31,35).

### Effects of the ITGB7, RFC5 and NOA1 on various traits

The effects of heterozygosity or homozygosity at the *ITGB7* substitution were estimated for 14 traits routinely recorded for evaluation purposes using the same model as for the haplotypes (see above; Supplementary Table 26), for age at first insemination and age at first calving (Supplementary Table 25), and for birth weight (Supplementary Table 30). Only the latter phenotype was analyzed for *RFC5* and *NOA1* due to insufficient numbers of homozygous mutant females with insemination and production record available.

### Estimation of the proportion of NOA1 homozygous mutants that died during embryonic development and between birth and genotyping

The conception rate (CR; i.e. the proportion of successful AI) was calculated for mating between males and females heterozygous for the NOA1 frameshift variant (1*1) and for mating between wild type parents (0*0) at two developmental stages (heifers and primiparous cows; Supplementary Table 32). The CR was expressed as a proportion of the CR observed in control mating for each female category (Centered_CR). The proportion of embryonic lethality among homozygous mutant conceptuses was calculated as (Centered_CR_0*0 - Centered_CR_1*1)/0.25, assuming that 25% of the conceptuses from at-risk mating are expected to be homozygous mutants, and full penetrance of embryonic lethality in the latter group. In parallel, we counted the proportion of each genotype observed in the offspring of heterozygous parental pairs and estimated the proportion of homozygous mutants genotyped out of the proportion expected according to Mendelian rules. These two values were used to derive the proportion of homozygous mutants that were born but died before reaching the age of genotyping (see Supplementary Table 33 for more information).

### Pathophysiological examination

*ITGB7*, *RFC5*, and *NOA1* homozygous mutant cattle aged two months to three years old and their matched controls were examined on farms. Blood samples were collected for serum and cell content analysis. Some of them were hospitalized for clinical follow-up at the bovine clinic of one of the four French National Veterinary Schools (Ecoles Nationales Vétérinaires de France, ENVF). During this time, routine blood analyses (hematology and biochemistry) and specific dosages (beta-hydroxybutyric acid, non-esterified fatty acids, lactate, etc.; see Supplementary Table 36) were performed at the central clinical pathology laboratory of each school. Euthanasia and necropsy were performed to identify gross lesions and to obtain tissue samples for further analysis to confirm hypothesized mechanisms of pathophysiology. Details on the number and age of animals considered are provided for each analysis in the figures in the Results section and in the Supplementary Tables.

### Hemogram

Blood was drawn from the jugular vein into 4-mL K 3-EDTA tubes (Venosafe, Terumo, France) and gently mixed by inversion. Tubes were immediately refrigerated at ∼4°C until hematologic analysis at the central laboratory of each school. Air-dried blood smears were prepared and stained with a May-Grünwald/Giemsa automated stainer (Aerospray hematology slide stainer cytocentrifuge 7150, Wescor, USA) for microscopic evaluation. Measurements were performed on a Sysmex XT-2000iV analyzer as recommended by the manufacturer, using settings for bovine blood (Sysmex XT-2000iV software v.00-13) (Herman et al. 2018). Analyzer-measured variables included red blood cell (RBC) count by impedance (RBC-I) and optical (RBC-O) measurements, hemoglobin concentration (HGB), and white blood cell (WBC) count. Neutrophil, lymphocyte, monocyte, eosinophil, and basophil counts were determined from 100 leukocytes counted per oil immersion field (1,000×). The percentages of each cell type were determined and the corresponding cell counts were calculated from the WBC.

### Metabolite analysis

Anticoagulated tubes were centrifuged within half an hour to prevent further exchange of analytes between blood cells and plasma. One mL of EDTA blood was mixed with 6% (w/v) perchloric acid for the determination of lactate. Biochemical analyses were performed on a Konelab 30 (Thermo Fisher Scientific inc., USA) using reagents from the same company, except for the determination of plasma lactate (Diasys Diagnostic Systems, Germany) or GLDH (Roche Diagnostic, Switzerland). Plasma troponin I concentration was determined by Immulite 2000 (Siemens, Germany). The HClO4 extracts were neutralized with 20% (w/v) KOH before analysis according to Baird and Heitzman 1970. Biological parameters were compared with reference samples prepared from healthy animals according to ASVCP recommendations (Friedrichs et al. 2012).

### Quantification of CD4+ mem a4+ b7+ T cells in the jejunal lamina propria using flow cytometry

One gram of small intestinal wall in the jejunal loop from heifers (n=3 homozygous mutant and 5 wild type) aged 1.5 to 3 years was washed in cold PBS, cut into 0.5 cm pieces, incubated four times in 30 mL of PBS 3 mM EDTA (Sigma-Aldrich), and digested in 20 mL of DMEM added with 20% FCS and 100 U/ml of collagenase (Sigma-Aldrich) for 40 min at 37 °C. Jejunum LP mononuclear cells were isolated on a 40–80% Percoll gradient after centrifugation at 1800 g for 15 min at room temperature. Then 1-2 x 10^6^ Mononuclear cells were resuspended in HBSS, 0.5% BSA, and 10 mM Hepes. Cell viability was assessed using Viobility 405/520 Fixable Dye (Miltenyi Biotec, Germany). Incubation with the antibodies was performed at 4°C for 30 min in the dark. The antibodies used were: CD4-PB (clone CC8, Biorad, Hercules, U.S), CD45-FITC (clone 1.11.32, Biorad), CD45R0-A647 (clone IL-A116, Biorad), Beta7-RPE (clone FIB27, Biolegend, San Diego, U.S.), Alpha4-PECy7 (clone 9F10, Biolegend). Data were collected on a MACSQuant® Analyzer (Miltenyi Biotec) and analyzed using Flowlogic software (Miltenyi Biotec).

### Light microscopy

Shoulder skin samples from four heifers homozygous for the *RFC5* inframe deletion and four matched controls, were fixed in 10% neutral buffered formalin for 24 h at +4°C, and then dehydrated in a graded ethanol series (30% to 100%), cleared in xylene, and embedded in paraffin wax. Longitudinal microtome sections (5 µm, Leica RM2245) were mounted on adhesive slides (Klinipath-KP-PRINTER ADHESIVES), deparaffinized, and stained with a Roan solution (nuclear fast red, orange G, and aniline blue). Slides were digitized with the Pannoramic Scan 150 and analyzed with CaseViewer 2.4 software (3D Histech). The number of hair follicles in a randomly selected 1 mm2 square and the diameter of the pilary canal of 50 adjacent hair follicles were measured for each animal (Supplementary Table 29).

### Transmission electron microscopy

Left ventricular heart samples from three heifers homozygous for the *NOA1* frameshift variant and three matched controls were fixed with 2% glutaraldehyde in 0.1 M Na cacodylate buffer pH 7.2, for 4 hours at room temperature. The specimens were then contrasted with Oolong Tea Extract (OTE) 0.2% in cacodylate buffer, postfixed with 1% osmium tetroxide containing 1.5% potassium cyanoferrate, dehydrated in a graded ethanol series (30% to 100%), and embedded in Epon, after the ethanol was gradually replaced by ethanol-Epon mixtures. Thin sections (70 nm) were collected on 200 mesh copper grids, and conterstained with lead citrate. The grids were examined with a Hitachi HT7700 electron microscope operated at 80kV (Milexia, France), and images were acquired with a charge-coupled device camera (AMT).

### Analysis of mitochondrial/nuclear DNA ratio by quantitative PCR

After preliminary testing focused on specificity and efficiency of amplification, two sets of mitochondrial and nuclear genes were selected with primer pairs showing similar slope (*CYTB* and *PPIA*, and *COX1* and *RSP24*, respectively; Supplementary Tables 37,38). Quantitative PCR was performed in triplicate on a QuantStudio 12K Flex Real-Time PCR System (Thermo Fisher Scientific) for five homozygous carriers of the *NOA1* frameshift insertion and five matched controls. The reaction mixture contained 10 μL 2X SYBR Green PCR Master Mix (Thermo Fisher Scientific), 1 ng total DNA extracted from myocardial samples, and 300 nM forward and reverse primers, in a total volume of 20 μL. For each animal the ΔCt(*CYTB*-*PPIA*) and ΔCt(*COX1*-*RSP24*) values were calculated based on the mean cycle threshold (Ct) of the triplicates. Finally the 2^-ΔΔCt(CYTB-PPIA)^ and 2^-ΔΔCt(COX1-RSP24)^ were used as two different indicators to measure the relative changes in the ratio of mitochondrial to nuclear DNA between the case and control groups.

## Ethics statement

All experiments reported in this work comply with the ethical guidelines of the French National Research Institute for Agriculture, Food and Environment (INRAE) and its French research partners. Blood samples from affected and unaffected animals were collected by licensed veterinarians. No animal was procreated purposedly for this study. The owners of the animals had consented to the diagnostic examination for research purposes and to the inclusion of their animals in this study. Invasive sampling was performed post-mortem only. As a consequence, no ethical approval was required for this study. Finally, all the data analyzed in the present study were obtained with the permission of breeders, breeding organizations and research group providers.

## Data Availability

Raw sequencing data from 732 cattle reported in this study have been deposited in the European Variation Archive (EVA, https://www.ebi.ac.uk/eva/) under accession numbers PRJEB64022 and PRJEB64023. Sequences from 1137 cattle from previous studies are available in the NCBI BioProject and EVA databases under the accession numbers listed in Supplementary Table 14. Source data for the other analyses are provided in the Supplementary Table file with this article.

## Acknowledgements

We are grateful to L. Balberini (Auriva), M. Chambrial (Origenplus), M. Courdier (Evajura), G. Fayolle (Umotest), C. Hamelin (Innoval), M. Philippe (Synetics), S. Patey (Genes Diffusion), N. Cesbron (Oniris), and the many breeders, veterinarians and agricultural technicians involved in this study for providing access to animals and samples. We also thank M. Bernard, C. Bevilacqua, E. Doz-Deblauwe, M. Femenia, C. Fouéré, N. Gaiani, J. Ros and N. Winter (INRAE) for their assistance and GenEval for providing data on yield deviations and genetic standard deviations. F.B. is supported by a CIFRE PhD grant from IDELE, with the financial support of the Association Nationale de la Recherche et de la Technologie and APIS-GENE (Paris, France). This study was also supported by the Effitness and Welcow projects funded by APIS-GENE, by the Cartoseq (ANR10-GENM-0018) and Bovano (ANR-14-CE19-0011) projects co-funded by the Agence Nationale de la Recherche and APIS-GENE, and by the SeqOccIn project co-funded by the European Union and the Occitania Region (FEDER-FSE MIDI-PYRENEES ET GARONNE 2014-2020).

## Author contributions

Conceived and coordinated the project: A. Capitan. Designed the experiments: A. Capitan, F.B., G.F., D.B. and M.-A.A. Performed HHED mapping: A.G. and A. Capitan. Set up haplotype tests: S.F. Analyzed survival curves: A.G., A. Capitan and C. Escouflaire. Analyzed pedigree information: F.B. and S. Minéry. Extracted DNA: C. Grohs and M.-C.D. Were involved in library preparation and whole genome sequencing: C. Eche and C.I. Processed WGS data: M. Boussaha, C. B., C. Klopp, M. Charles and C. Kuchly. Filtered and annotated variants: F.B., A. Capitan and A.G. Analyzed large-scale genotyping data: F.B., A. Capitan and C. Escouflaire. Estimated the effect of haplotypes or deleterious variants on a variety of traits: F.B., A. Capitan, J.J. and A. Barbat. Performed the pathophysiological analyses and necropsy examinations: G.F., L.G.-P., M.-A.A., A. Clément., L.D., B.G., E.C., M. Bouchier, T.B., A. Remot, V.P., A. Relun, B.R., Y.M., R.G. Assisted in sampling: A. Capitan, C. Grohs, M-C.D., M. Cano. Performed histological and ultrastructure analyses: M. Cano, C.P., A. Capitan, J.R. and M.V. Performed qPCR analysis: C. Grohs. Contributed reagents/materials/analysis tools: C.H., H.L., A. Boulling, S.B., C. Grohs, C.D.-B., F.L., S. Mattalia, A.A.B., G.V., C. Gaspin, C.D. and D.M. Wrote the manuscript: A. Capitan and F.B. with contributions from G.F., L.G.-P., D.B. J.J. and M.-A.A.

