## Supplementary Figure 1 for "Massive detection of cryptic recessive genetic defects in cattle mining millions of life histories"

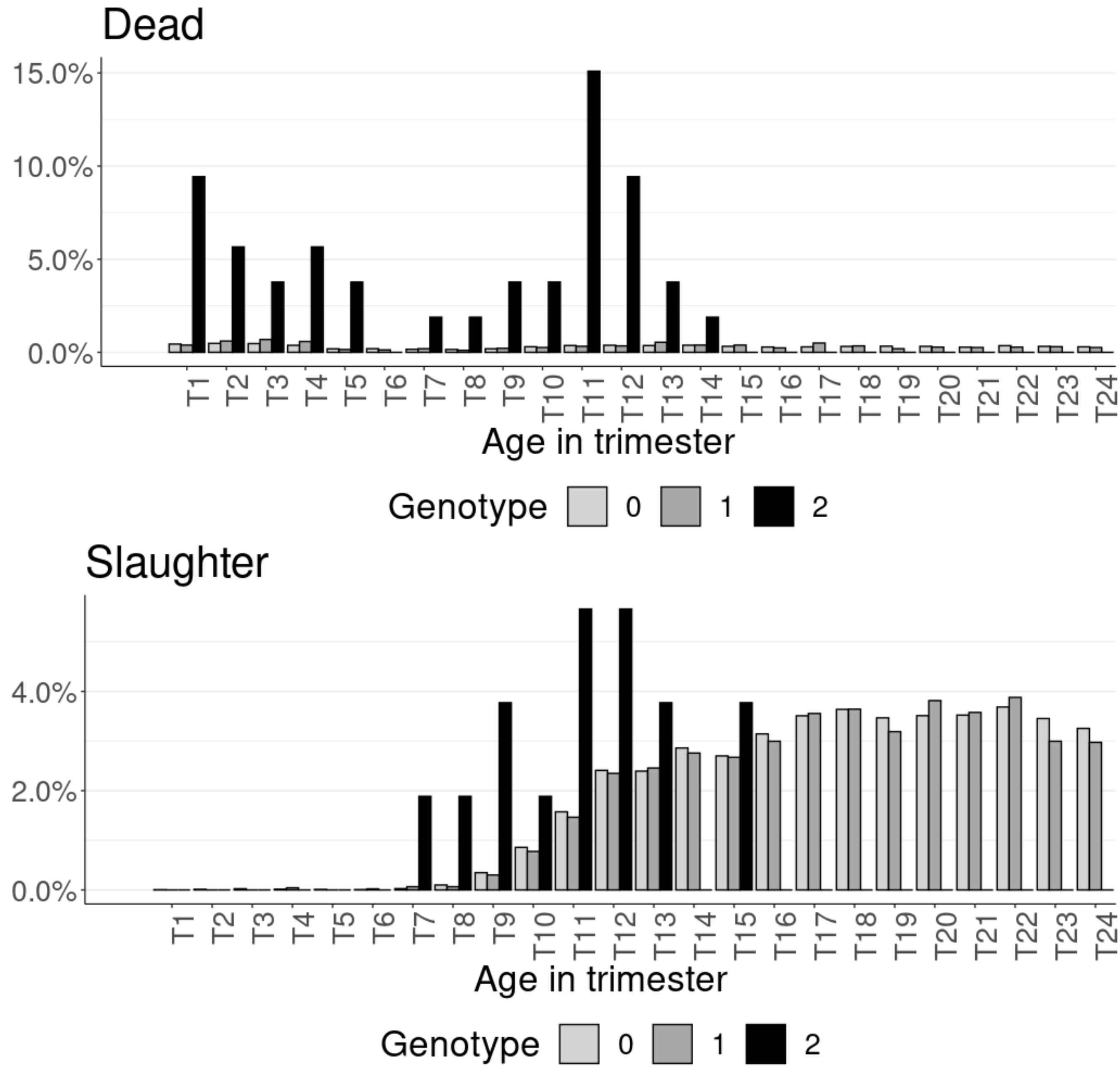


**Supplementary Figure 1. Trimester repartition of the proportion of females that died or were slaughtered before reaching six years of age by genotype group at the *RFC5* substitution.** Source data are provided in Supplementary Table 31.
