## Supplementary Figure 2 for "Massive detection of cryptic recessive genetic defects in cattle mining millions of life histories"

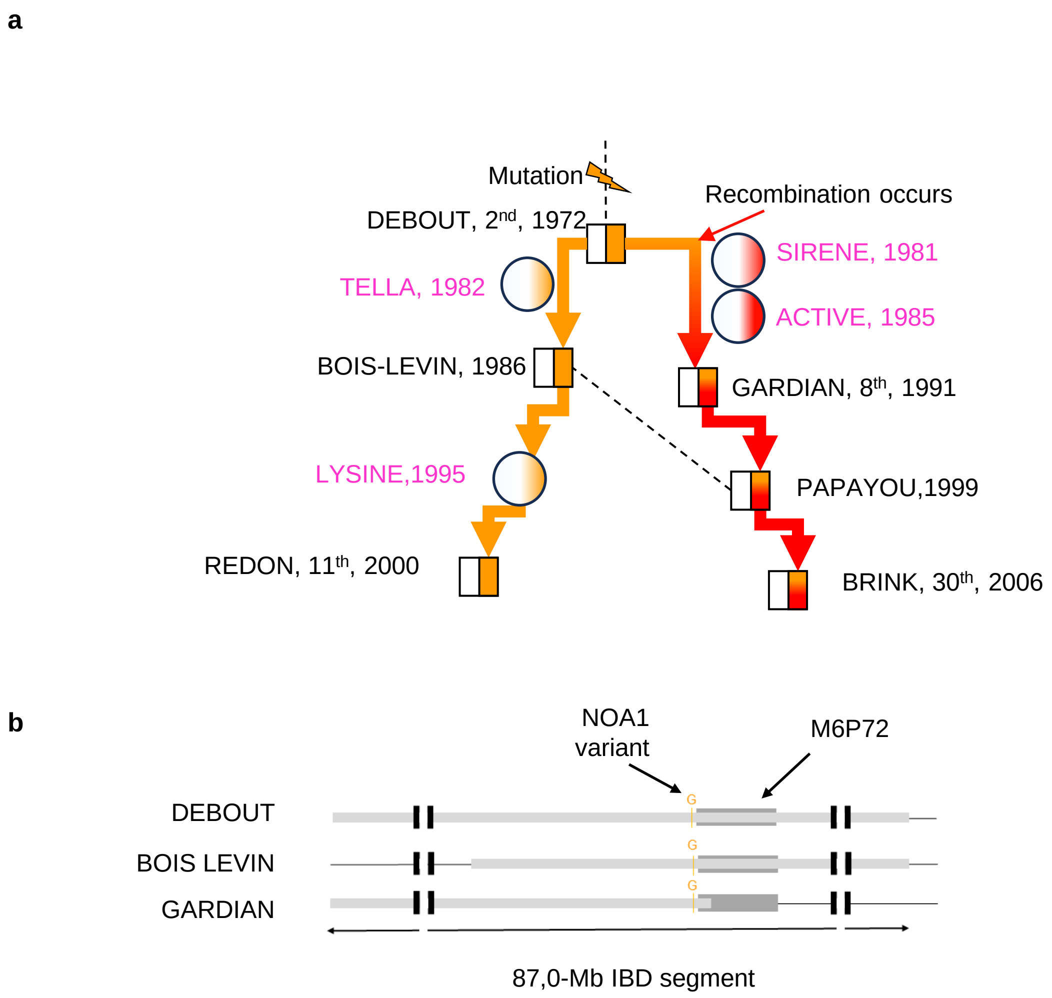


**Supplementary Figure 2. Analysing the origin of M6P72 haplotype and NOA1 frameshift variant.** a) Schematic pedigree showing the dissemination of the haplotype and NOA1 variant following a founder effect and subsequent recombination events in the French Montbeliarde breed. The 2nd, 8th, 11th, and 30th largest contributors to the breed are highlighted (Supplementary Table 12). Several chromosomal segments carry the mutation but only one is detected by the haplotype test. b) Details of the IBD segments shared by the bulls DEBOUT, BOIS-LEVIN and GARDIAN showing the recombination in the middle of the M6P72 haplotype.
