## Supplementary Note for "Massive detection of cryptic recessive genetic defects in cattle mining millions of life histories"

### Supplementary Note. Neutrophils from *NOA1* homozygous mutant calves are functional

#### Results

Since Nitric Oxide-Associated Protein 1 (NOA1) has been shown to play a role in plant immunity (e.g., Cobessi et al., 2012; Frederickson Matika and Loake, 2014) and nitric oxide (NO) synthesis is involved in the antibacterial immune response in animals (e.g., MacMicking et al., 1997; Lundberg and Weitzberg, 2022; Guzik et al., 2003) we examined the production of reactive oxygen species (ROS) by blood neutrophils and monocytes from calves homozygous for a NOA1 frameshift mutation and matched controls by flow cytometry (**Supplementary Note Fig. 1A**). The ROS production by neutrophils (G1+) or monocytes (CD14+) was similar between cases and controls, both at basal level and upon stimulation with tert-butyl hydroperoxide (**Supplementary Note Fig. 1B**). We also measured the specific levels of $H_2O_2$ and NO in neutrophils by spectrophotometry and again observed no obvious difference between cases and controls (**Supplementary Note Fig. 1C**). We concluded that ROS production by monocytes and neutrophils was not affected by the NOA1 mutation.

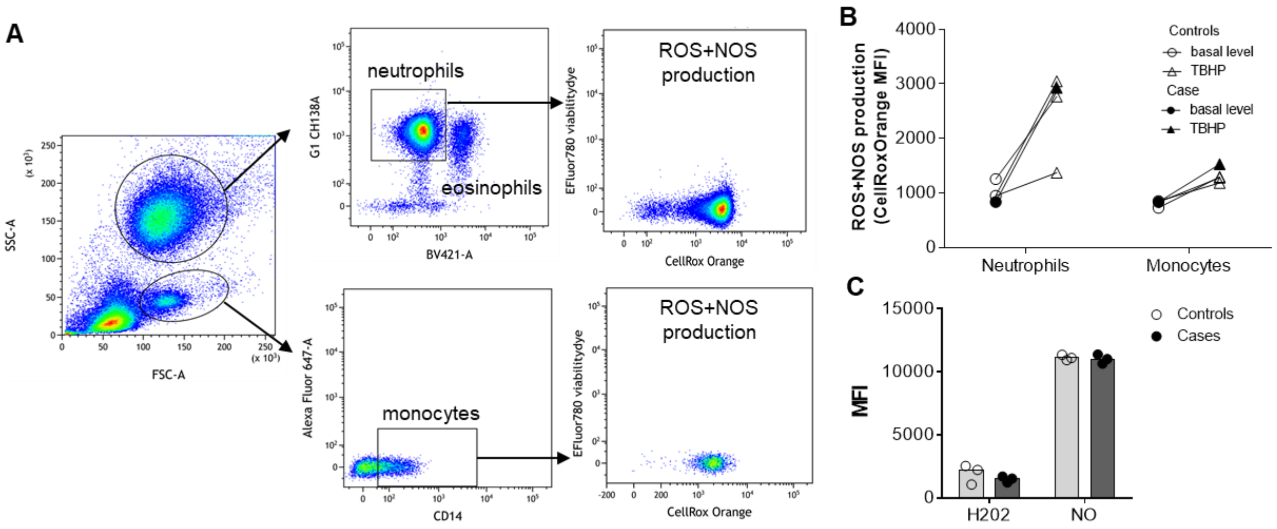


**Supplementary Note Figure 1. Neutrophil oxidative stress levels are similar between cases and controls.** A) Flow cytometry gating strategy used to identify blood neutrophils (granulocytes autofluoBV421- G1+) and monocytes (CD14+). ROS and RNS production were measured by flow cytometry using the cellRoxOrange dye. B) Mean Fluorescence Intensity (MFI) of the cellRoxOrange dye in the neutrophils or monocytes gates, measured with or without 1h ex vivo stimulation with tert-butyl hydroperoxide TBHP (circles and triangles, respectively). Results are shown for three controls (clear symbols) and one case calf (black symbols; representative of 2). C) MFI of the H2DCFDA and DAF2DA dyes (H2O2 and NO respectively) produced ex vivo by neutrophils (positive enrichment with G1 Ab and magnetic beads) measured with the TECAN SparkControl. Results are shown for three controls (clear symbols) and three cases (black symbols).

We then extracted RNA from case and control neutrophils, to evaluate the transcription of key neutrophil enzyme genes: myeloperoxidase, elastase, and cathepsin G, and the pro-inflammatory cytokine *IL-1b*. As shown in **Supplementary Note Fig. 2**, gene expression was equivalent in the two groups, suggesting that neutrophils from cases are properly equipped to respond to infection.

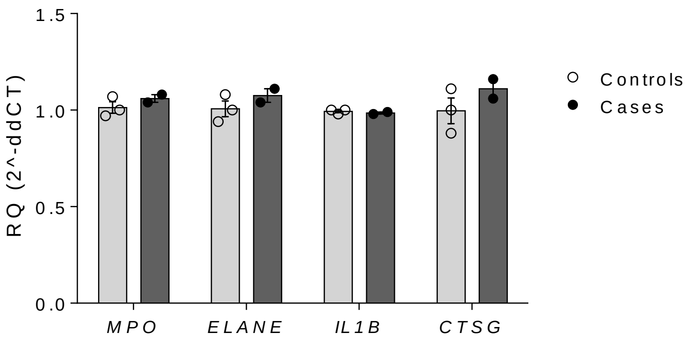


**Supplementary Note Figure 2. Gene expression in neutrophils.** Neutrophils were enriched by positive magnetic bead sorting (G1 Ab) and RNA was extracted. The expression of 4 neutrophil-related genes, myeloperoxidase (*MPO*), elastase (*ELANE*), IL-1b (*IL1B*) and cathepsin G (*CTSG*) was measured by qRT-PCR, and normalized to the expression of 3 housekeeping genes (*PPIA*, *ACTB,* and *GAPDH*) and then to the control group (ddCT). Individual data of 2 case and 3 control animals are presented.

In conclusion, the comprehensive analysis performed did not reveal any discernible differences between the two groups in the parameters evaluated. Therefore, no inherent defects in neutrophil function could be demonstrated in this study.

#### Methods

##### Preparation and labeling of neutrophils

Blood samples were collected by veterinarians and sent to INRAE Nouzilly. Upon receipt 18-24 hours after sampling, Vacutainer K2 EDTA tubes (10 ml) were centrifuged at 1,000g for 15 min at 20°C. Plasma and buffy coat were discarded, then ACK (Gibco) (5 vol/1 vol blood cells) was added for 5 min at room temperature to lyse red blood cells. Cells were washed twice in D-PBS (without calcium and magnesium) with 2mM EDTA, resuspended in D-PBS (without Ca/Mg) containing 10% of horse normal serum (Gibco) and 2mM EDTA, and labeled for 30 min at 4°C with primary antibodies: mouse anti-bovine granulocytes G1 (CH138A). After washes in D-PBS with 2mM EDTA and 10% of horse normal serum (400g, 10 min, 4°C), cells were labeled with the appropriate fluorescent-conjugated secondary antibodies (goat anti-mouse IgM A647, goat anti-mouse IgG1 A488, and the eFluor780 Fixable Viability Dye) for 30 min at 4°C. Cells were washed and i) plated for ROS/NOS assay or ii) analyzed by flow cytometry by direct acquisition using an LSR Fortessa™ X-20 flow cytometer (Becton Dickinson) or iii) purified as described below.

##### Neutrophil purification

Cell concentrations were adjusted to $10^7$cells/mL and sorted with a MoFlo AstriosEQ high-speed cell sorter (Beckman Coulter) or with anti-G1 Ab and anti-IgM magnetic beads according to our previously published protocol (Rambault et al., 2021). Sorted cells were used for RNA preparation or spread on microscope slides (Superfrost, Thermo) by cytocentrifugation (3 min, 700xg) and stained with May-Grünwald and Giemsa using the RAL 119 555 kit (RAL Diagnostics).

##### ROS/NOS Assay

ROS/NOS production was quantified using the CellROX® Orange Flow Cytometry Assay Kits (MolecularProbes®, C10493) according to the manufacturer's instructions. Briefly, $10^5$ blood cells were first incubated for 1 hour at 37°C in RPMI supplemented with 1 mg/mL BSA, 1 mM EDTA, and 10 mM HEPES with or without 400 µM of TBHP in a black 96-well microplate and then for 30 min with 100 nM CellROX® and 1 µL/mL efluor780 fixable viability dye. Fluorescence was measured directly (without fixation) using the LSR Fortessa™ X-20 flow cytometer. Flow cytometry results were analyzed using Kaluza software (Beckman Coulter).

Alternatively, H2O2 and NO production were assessed on plated cells with $H_2$DCFDA and DAF$_2$DA dyes, respectively, according to the manufacturer's instructions (Sigma). Fluorescence was measured using the TECAN SparkControl and the Magellan software.

##### Gene expression

Total RNA was extracted using a NucleoSpin RNA kit with a DNase treatment (Macherey Nagel) and reverse transcribed with iScriptTM Reverse Transcriptase mix (Biorad) according to the manufacturer's instructions. qRT-PCR were performed with a LightCycler® 480 Real-Time PCR System (Roche), with the primers (Eurogenetec) listed in the supplementary Table1. The annealing temperature was 60°C. Data were analyzed with LC480 software to determine the cycle threshold (Ct) values. Messenger RNA expression was normalized to the mean expression of three housekeeping genes to obtain the ΔCt value. For each gene, values from cases were normalized to the mean expression of the control group (ΔΔCt value, and Relative Quantity = $2^{-\Delta\Delta Ct}$).

| Gene symbol | Full name | Primer pairs |
|---|---|---|
| *GAPDH** | *Glyceraldehyde-3-phosphate dehydrogenase* | GGCATCGTGGAGGGACTTATG<br>GCCAGTGAGCTTCCCGTTGAG |
| *ACTB** | *Actin beta* | ACGGGCAGGTCATCACCATC<br>AGCACCGTGTTGGCGTAGAG |
| *PPIA** | *Peptidylprolyl isomerase A* | TCCGGGATTTATGTGCCAGGG<br>GCTTGCCATCCAACCACTCAG |
| *MPO* | *Myeloperoxidase* | AATGACCCCCGCATCAAGAA<br>CGTTGATCTGGTTGCGGATG |
| *ELANE* | *Elastase* | CGATTCCTTCATCCGTGGGG<br>GCGCCGGATGATAGAGTTGA |
| *IL1B* | *Interleukin 1 beta* | CTCTCACAGGAAATGAACCGAG<br>GCTGCAGGGTGGGCGTATCACC |
| *CTSG* | *Cathepsin G* | ATTTCCAGCTTCCTGCCCTG<br>TGCCCCAGAGAAGGAGAGTC |

**Supplementary Note Table 1. Primers used for qRT-PCR.** *: Housekeeping genes.
